# Prevalence and Determinants of Caesarean Section in South and South-east Asian Women

**DOI:** 10.1101/616813

**Authors:** Vivek Verma, Ramesh K. Vishwakarma, Dilip C. Nath, Omer Abid, Ram Prakash

**Affiliations:** AIIMS Cohort Study Centre, All India Institute of Medical Sciences, New Delhi, India; Department of Biostatistics, King Abdullah International Medical Research Center, Riyadh, Saudi Arabia; King Saud bin Abdulaziz University for Health Sciences, Riyadh, Saudi Arabia; Ministry of the National Guard - Health Affairs, City, Saudi Arabia; Assam University, Silchar, Assam, India; Department of Population Health Research, King Abdullah International Medical Research Center, Riyadh, Saudi Arabia; King Saud bin Abdulaziz University for Health Sciences, Riyadh, Saudi Arabia; Ministry of the National Guard - Health Affairs, City, Saudi Arabia; Department of Quality Assurance Unit, Eurofins Therapeutics Limited, Bangalore, India

## Abstract

**Background:** Caesarean section is considered a preferable and safe method of delivery. In the last decade, its prevalence has increased in both developed and developing countries. In the context of developing countries viz., South Asia (the highest populated region) and South-east Asia (the third highest populated region), the preference for, and variation in, caesarean section delivery and its associated maternal socioeconomic characteristics are still to be determined.

**Objective:** To study the magnitude of caesarean delivery in the South and South-east Asian countries, by correlating the maternal socioeconomic characteristics with the preference for caesarean sections.

**Methodology:** Data on ever-married women of nine developing countries of South and South-east Asia *viz.*, Vietnam, India, Maldives, Timor-Leste, Nepal, Indonesia, Pakistan, Bangladesh and Cambodia, from Demographic and Health Survey has considered. Both bivariate and multivariate binary logistic regression models were used to estimate the probability that a woman undergoes caesarean section and to assess the influence of maternal socioeconomic characteristics towards the preference for caesarean section.

**Conclusion:** In seven urban and four rural regions of nine South and South-East Asian countries, a significant inclination towards the caesarean delivery above the more recent outdate WHO recommended optimal range of 10-15% or the more recent study by top researchers of 19% has been found. The analysis confirmed that the prevalence of caesarean section and its associated maternal socioeconomic characteristics varied widely among these nine South and South-East Asian countries.

## Introduction

Caesarean section in developed and developing countries [1]–[3], has considered as the most preferred method of child birth. The preferences of caesarean section is found to be comparatively high in the last decade [4], and one of the significant reason for this is the reduction in risk of death to mother and child during delivery [5], [6]. The reasons for the increase in C-section are due to various factors. In most of the developing countries, demographic changes, social and educational improvement have led to an growing number of women delaying their pregnancies until getting on their end of fertile life [7]. This social development pooled with the approachability to birth control and infertility treatment has increased the number of women experiencing their first pregnancy only after 35 years of age [8]. The Caesarean section or C-section is a surgical procedure, where delivery proceeds through the abdominal and uterine incision. This procedure is appropriate in situations where vaginal (or normal) delivery will increase life risks for the mother and the baby. Although caesarean delivery is considered a relatively safe delivery method, the risk of complication is higher [9] as compared to a vaginal birth or a natural method of birth. One of the major issue with the caesarean deliveries other than the post-delivery risks and complication is the cost, which increases due to the operation and longer stays in hospitals, and that creates a huge financial burden to the family [10]. The most frequently occurring complications during and after a caesarean to mothers and children have already been discussed [11]–[14]. In the past, the World Health Organization [15], had suggested that although caesarean section is a safe method, but if caesarean rate exceeds the limit of 10-15%, it may not lead to better outcomes. However, that previous suggestion had come under criticism for multiple reasons. The WHO may have changed it’s view as it released a statement in 2015 with the headline. Every effort should be made to provide caesarean sections to women in need rather than to achieve a specific rate. Earlier works [16]–[18] have suggested that if caesarean rate increases above WHO recommended range, then as a consequence the risk of manifestation of other public health-related problems of both mothers and children will also increase. Some of work [19] has more recently concluded that the 1985 WHO document [15] looked at studies that were incomplete because they examined data from limited sets of countries and often examined outcomes in wealthier countries. In addition, many studies used data from varying years without accounting for heterogeneity across years.

Furthermore, what is being overlooked is that the WHO document [15] looked at only a correlation only with mortality. Morbidity, both fetal and maternal, were not taken into account for these rates. It is essential to keep foremost in mind that fetal morbidity should be weighed much more higher than maternal morbidity as failure to do C-section when indicated can result in babies with profound brain damage which are catastrophic not only for the babies entire future life but also catastrophic for the parental caregivers and the rest of the family.

The rate of caesarean section is usually defined as the fraction of women who adopted a caesarean delivery procedure among total childbirths in a specified time-period in a specific geographic area”. The prevailing models and estimates of the caesarean rate in a specific geographic area are appropriate under the assumption that in this selected area almost all deliveries took place in medical institutions, as the procedure of caesarean delivery is possible only if delivery is institutional. But in developing countries of South and South-east Asia *viz.*, Vietnam, India, Maldives, Timor-Leste, Nepal, Indonesia, Pakistan, Bangladesh and Cambodia, a considerable proportion of child deliveries are carried out at home (Table 1) and are completely free of risks and complications associated with caesarean deliveries.

As the caesarean is a surgical procedure and is only possible at medical institutions; Therefore, in the present study, the prevalence of caesarean section among women from different South and South-east Asian countries, who have experienced the institutional deliveries, have been investigated. In this study, an attempt has been made to provide a better understanding of the behavioural pattern of among women of these countries towards caesarean section. Through this study, the dependency and importance of the socio-economic factors on the caesarean section preference for delivery have also been explored.

### Socio-Economic Factors Affecting C-Section procedures

The countries of South and South-east Asia are regions of great social, economic and political diversity. Despite their diversity, countries in this region are attempting to improvise their regional health challenges ([20] and [21]) and promoting safe and healthy maternal and child health, and also encourage deliveries in presence of trained health professionals [22], under proper hygienic conditions, as the caesarean procedure is only possible at a medical institution.

The relationship with incidences of caesarean section with some of associated maternal socio-economic characteristics *viz.*, maternal age, place of residence, level of maternal education, birth order of the child, and type of medical facility that opted for delivery and size of the baby, has been modelled. Age as reported subjectively by mother, and grouped into 6 subgroups: 15-19, …, 40-44. The 45 to 49 age group was not taken into account due to a lack of sufficient data. The type of place of residence has classified as rural and urban. The educational qualification has classified into four classes, primary, secondary and higher. Birth order of the born child, which has grouped into first, second, third, fourth and fifth or more. The medical institutions that opted for the delivery were grouped together whether government or private facilities. The size of the child was classified as below average, average and above average and large. Our interest lies in to find the prevalence of caesarean section in the countries of South and South-east Asia.

## Methodology

### Data

Data for this study is obtained from Demographic and Health Surveys (DHS) database on maternal deliveries occurred in nine developing countries *viz.*, Vietnam, India, Maldives, Timor-Leste, Nepal, Indonesia, Pakistan, Bangladesh and Cambodia. DHS are series of nationally representative household surveys that provide information on population, health and nutritional status of mother and child. The study dataset includes only the latest round of data of each selected country. List of selected South and South-east Asian countries and the corresponding survey years are given in Table 1.

### Statistical analyses

Both bivariate logistic regressions and multivariate logistic regression models have constructed separately to estimate the prevalence of caesarean section to a woman based on her socio-economic characteristics, viz., maternal age, place of residence, level of maternal education, birth order of the child, and type of medical facility that opted for delivery and size of the baby. The results obtained from the regression analyses have been presented in terms of the odds ratios (ORs) with 95% confidence interval (CI).

**Table 1.** Selected DHS countries, survey years and place of birth of selected **South and South-east Asian Countries**

| S.No. | Country | Survey Year | Total Births* | Place of Birth |  |
| --- | --- | --- | --- | --- | --- |
|  |  |  |  | House (%) | Medical(%) |
| 1 | Vietnam | 2002 | 1317 | 20.20 | 79.80 |
| 2 | Maldives | 2009 | 3817 | 4.06 | 95.94 |
| 3 | Timor-Leste | 2009-10 | 9806 | 80.79 | 19.21 |
| 4 | Nepal | 2011 | 5306 | 62.34 | 37.66 |
| 5 | Indonesia | 2012 | 18021 | 45.08 | 54.92 |
| 6 | Pakistan | 2012-13 | 11763 | 50.24 | 49.76 |
| 7 | Bangladesh | 2014 | 7886 | 76.48 | 23.52 |
| 8 | Cambodia | 2014 | 7165 | 17.85 | 82.15 |
| 9 | India | 2015-16 | 259627 | 24.51 | 75.49 |

\*Total births occurred in last five year preceding the survey

Statistical analyses were performed using the Statistical Analysis System (SAS) package, (university edition) and all other computation is carried out using R (version-3.0.3) and SAS version 9.4. Corresponding to each of the associated maternal socio-economic characteristics, the associations with the prevalence of caesarean sections have been examined using binary logistic regression analyses, to examine the effect of socio-economic factors on the odds of caesarean birth and non-caesarean birth. The event of caesarean section during delivery, as a dichotomous variable, where if delivery is caesarean then denotes‘1’ delivery and ‘0’ for other than caesarean

## Results

### Background Characteristics of Prevalence of Caesarean Section

Table 2 presents the rate of caesarean section based on both institutional and non-institutional births, separated by place of residences *viz.*, rural and urban regions, of South and South-east Asian Countries. A substantial inter-region variation in caesarean rates corresponding to each country has been observed. Obtained results have shown an inclination of caesarean delivery among urban than rural women and are quite conspicuous. The rate of caesarean delivery found in urban part is highest in women of Maldives (39.07%) followed by India (23.64%), Bangladesh (21.82%), Vietnam (21.72%), Pakistan (17.75%) and Indonesia(17.25%), which have crossed the WHO recommended range of 10-15%, which might be better to have upper limit of 19% [19]. The caesarean rate in the rural women of Maldives (30.70%) has only crossed the WHO recommended range. The caesarean rate in women residing in urban regions is five times of rural women in Nepal; three times of rural women of Vietnam, Timor-Leste, Indonesia and Cambodia, and twice of the rural women of India, Pakistan and Bangladesh.

To obtain a better estimate for the prevalence of caesarean deliveries in each of the selected South and South-east Asian Countries, the population is classified into two disjointed sub-populations based on their place of delivery *viz.*, institutional or non-institutional. Here, institutional deliveries referred to those deliveries occurred at any private or government medical institutions, whereas births occurred other than any medical institutions, are considered as non-institutional births. To visualise the prevalence of caesarean section and the population at risk, only institutional deliveries have been taken into account for investigation.

**Table 2.** Rate of caesarean births occurred based on **both institutional and non-institutional births** in different **South and South-east Asian Countries** during the five years preceding the survey

| S.No. | Country | Urban |  | Rural |  | All |  |
| --- | --- | --- | --- | --- | --- | --- | --- |
|  |  | Births | Rate | Births | Rate | Births | Rate |
| 1 | Vietnam | 267 | 21.72 | 1050 | 7.05 | 1317 | 10.02 |
| 2 | Maldives | 494 | 39.07 | 3323 | 30.70 | 3817 | 31.78 |
| 3 | Timor-Leste | 2204 | 3.18 | 7602 | 1.03 | 9806 | 1.51 |
| 4 | Nepal | 1091 | 11.64 | 4215 | 2.78 | 5306 | 4.60 |
| 5 | Indonesia | 8170 | 17.25 | 9851 | 6.96 | 18021 | 11.63 |
| 6 | Pakistan | 4970 | 17.75 | 6793 | 8.05 | 11763 | 12.15 |
| 7 | Bangladesh | 2488 | 21.82 | 5398 | 10.10 | 7886 | 13.80 |
| 8 | Cambodia | 1950 | 12.97 | 5215 | 4.31 | 7165 | 6.67 |
| 9 | India | 61379 | 23.64 | 198248 | 10.68 | 259627 | 13.74 |

**Table 3.** Rate of caesarean births occurred based on **institutional births** in different **South and South-east Asian Countries** during the five years preceding the survey

| S.No. | Country | Urban |  | Rural |  | All |  |
| --- | --- | --- | --- | --- | --- | --- | --- |
|  |  | Births | Rate | Births | Rate | Births | Rate |
| 1 | Vietnam | 263 | 22.05 | 788 | 9.01 | 1051 | 12.27 |
| 2 | Maldives | 490 | 39.18 | 3167 | 32.21 | 3657 | 33.14 |
| 3 | Timor-Leste | 941 | 7.44 | 943 | 8.27 | 1884 | 7.86 |
| 4 | Nepal | 727 | 17.47 | 1271 | 9.21 | 1998 | 12.21 |
| 5 | Indonesia | 6246 | 22.49 | 3634 | 18.79 | 9880 | 21.13 |
| 6 | Pakistan | 3174 | 27.79 | 2674 | 20.46 | 5848 | 24.44 |
| 7 | Bangladesh | 862 | 62.88 | 993 | 54.78 | 1855 | 58.54 |
| 8 | Cambodia | 1846 | 13.71 | 4039 | 5.57 | 5885 | 8.12 |
| 9 | India | 53200 | 27.28 | 142797 | 14.82 | 195997 | 18.20 |

Results based on institutional delivery, Table 3 shows that the rate of caesarean delivery found is highest among the urban women of Bangladesh (62.88%) followed by Maldives (39.18%), Pakistan (27.79%), India (27.28%), Indonesia(22.49%), Vietnam (22.05%), and Nepal (17.47%) that have crossed the WHO recommended range of 10-15%. Among the rural women, the caesarean rate is on higher side of Bangladesh (54.78%) followed by Maldives (32.21%), Pakistan (20.46%) and Indonesia (18.79%), which have crossed the WHO recommended range. The overall caesarean rate in women of five countries *viz.*, India, Maldives, Indonesia, Pakistan and Vietnam, have found more than 15%.

The percentage distribution of caesarean section among women based on their socio-demographic characteristics have depicted in Figure 1–6. The pie-chart depicted in Figure 1 is showing a positive association of caesarean rate with the 15-39 age-grouped women, but on the contrary the caesarean rate among women belongs to 40-44 is comparatively low as compared to those belongs to 35-39. It also shows that the caesarean rates are more than 15% (WHO recommended) among the women aged 30 and above in all countries except Timor-Leste and Cambodia. Figure 2 depicts a negative association between the prevalence of caesarean delivery and the birth order of the child, *i.e.*, chance to get caesarean section is high to women having least birth order. In countries Maldives and Bangladesh, caesarean rates are very high and found for all birth orders. From Figure 3, it has found that caesarean is more preferred to women whose baby sizes are either very large or smaller than average. Irrespective of the size of the baby, caesarean rates in Maldives, Nepal and Bangladesh have been found to be very high. Figure 4 depicts a positive association between caesarean section and women’s education level. It is found in all countries that women who have a higher level of education are more predisposed towards caesarean childbirth. Caesarean sections among urban women, as depicted in Figure 5, are very high as compared to those of rural women except in Bangladesh, where rates are very high in women from both regions. Figure 6, depicts the caesarean sections among the women whose delivery occurred at private medical institutions which are very high compared to those delivered at government medical institutions.

**Fig 1.**
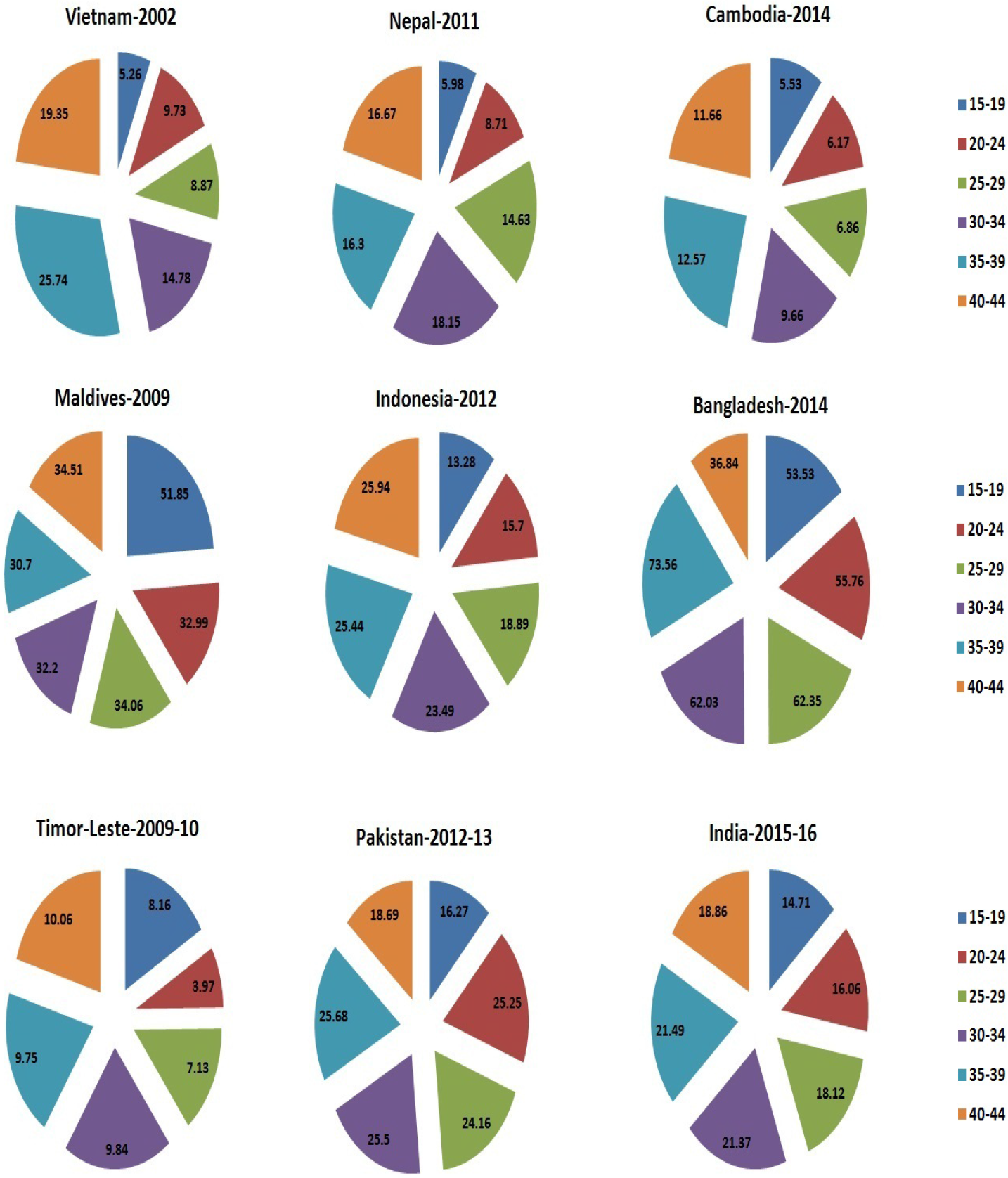
Caesarean rate based on mothers’ age-interval in South and South-east Asian Countries.

**Fig 2.**
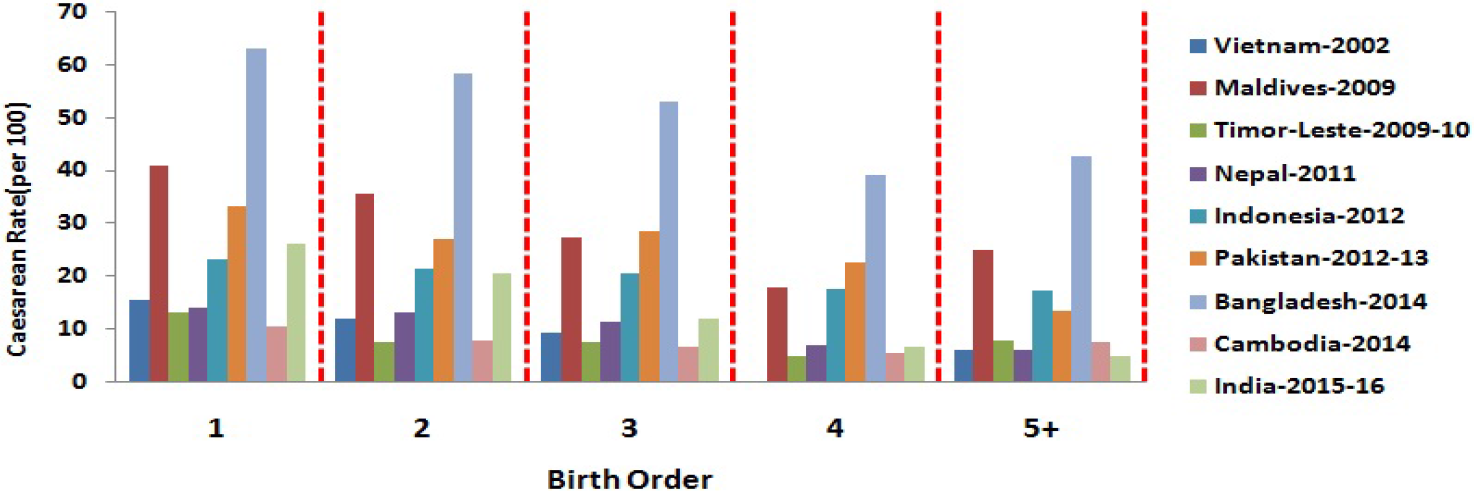
Caesarean rate based on birth order of the child in South and South-east Asian Countries.

**Fig 3.**
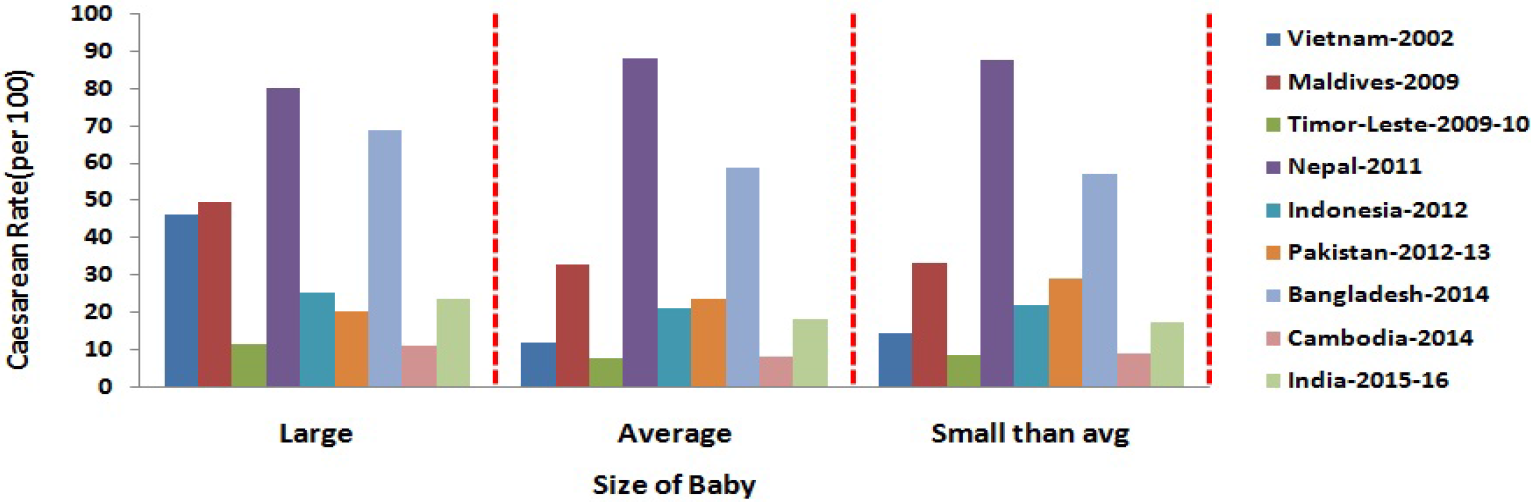
Caesarean rate based on size of the delivered child in South and South-east Asian Countries.

**Fig 4.**
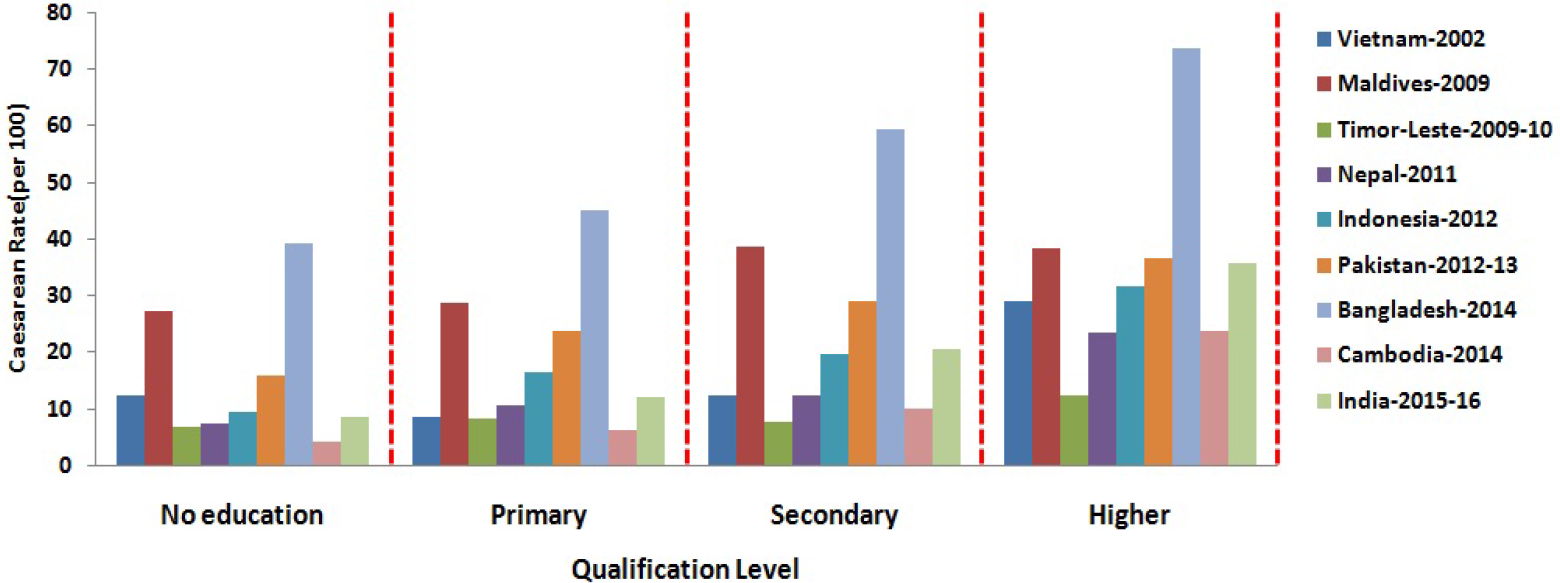
Caesarean rate based on mothers’ qualification in South and South-east Asian Countries.

**Fig 5.**
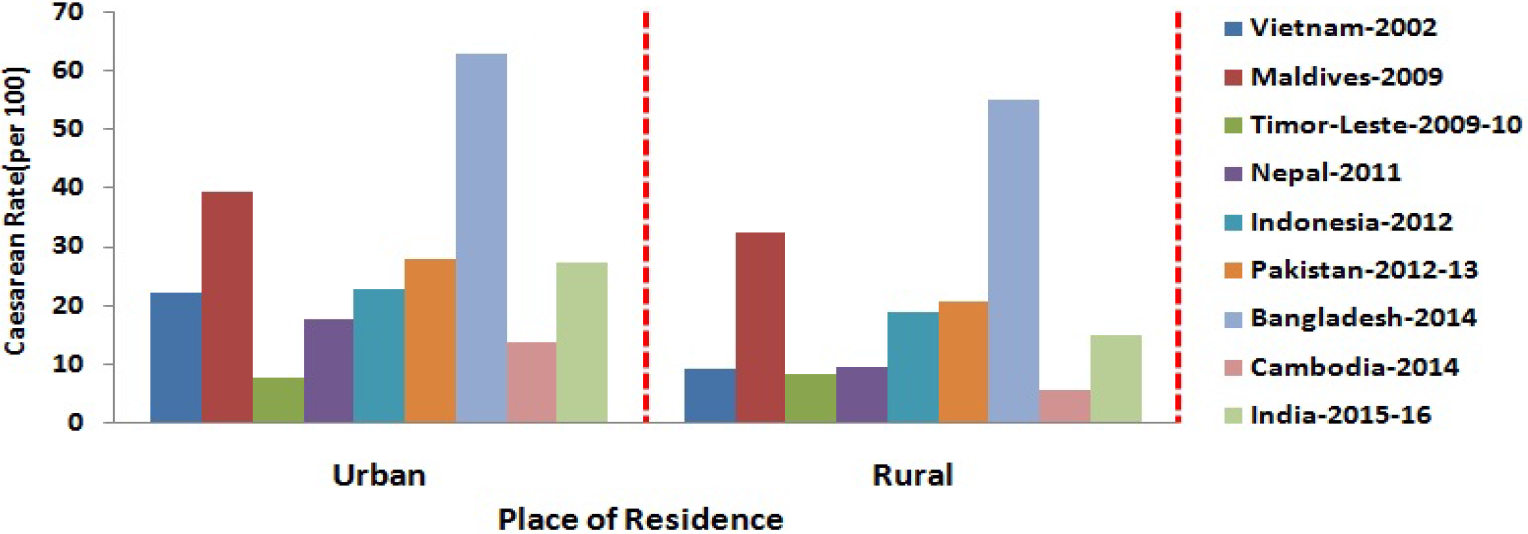
Caesarean rate based on mothers’ place of residence in South and South-east Asian Countries.

**Fig 6.**
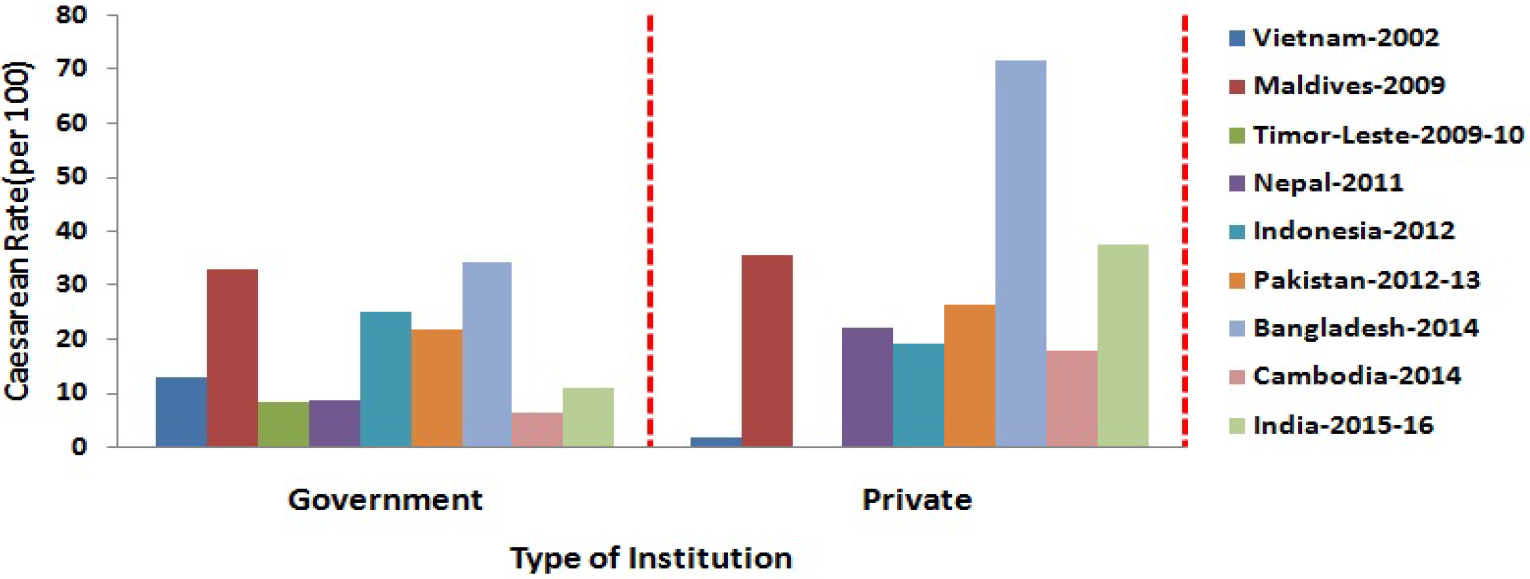
Caesarean rate based on choice of institution in South and South-east Asian Countries.

### Socio-Economic Determinants of Prevalence of Caesarean Section

Tables 4 and 5 shows the results of the bivariate and multivariate logistic regression analysis, respectively. The unadjusted analysis of multivariate logistic regression has revealed that maternal age, mother’s education (except Vietnam and Maldives), choice of medical institutions (government and private) for childbirth (except Vietnam and the Maldives), birth order and place of residence (except the Maldives, Timor-Leste and Bangladesh) have a significant effect on caesarean section. The variable baby size” has a significant effect on the prevalence of caesarean delivery (except Timor-Leste, Nepal, Indonesia, Bangladesh and Cambodia). The bivariate analyses applied in the study showed that maternal age, mother’s education (except Timor-Leste), choice of medical facility for childbirth (except the Maldives), birth order and place of residence (except the Maldives) have a significant effect on the prevalence of caesarean section. The variable ‘size of the baby’ had found to be insignificant in Timor-Leste, Nepal, Indonesia, Bangladesh and Cambodia, while in the remaining countries *viz.*,Vietnam, Maldives, Pakistan and India revealed has a significant effect on the prevalence of caesarean delivery.

**Table 4.**
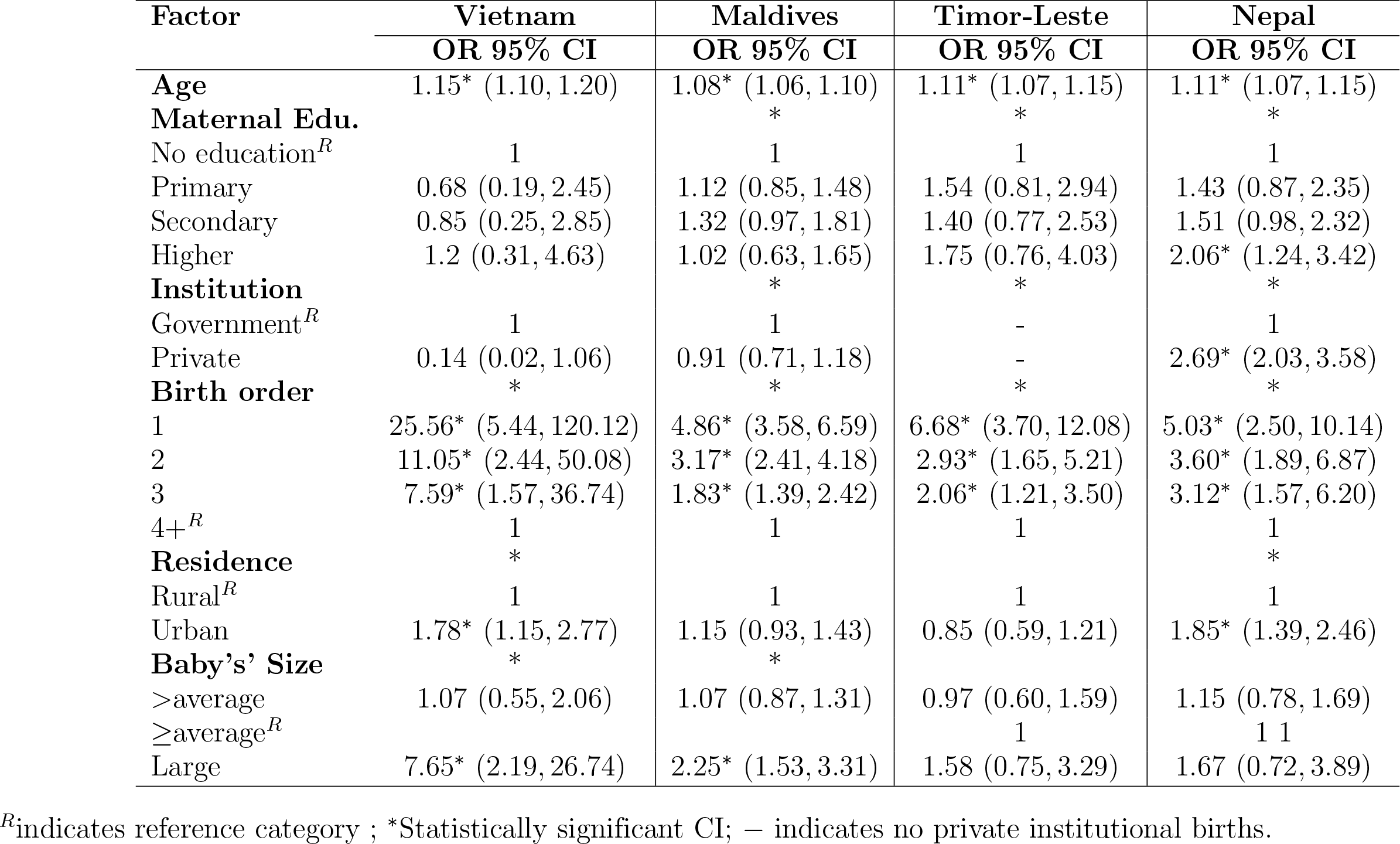
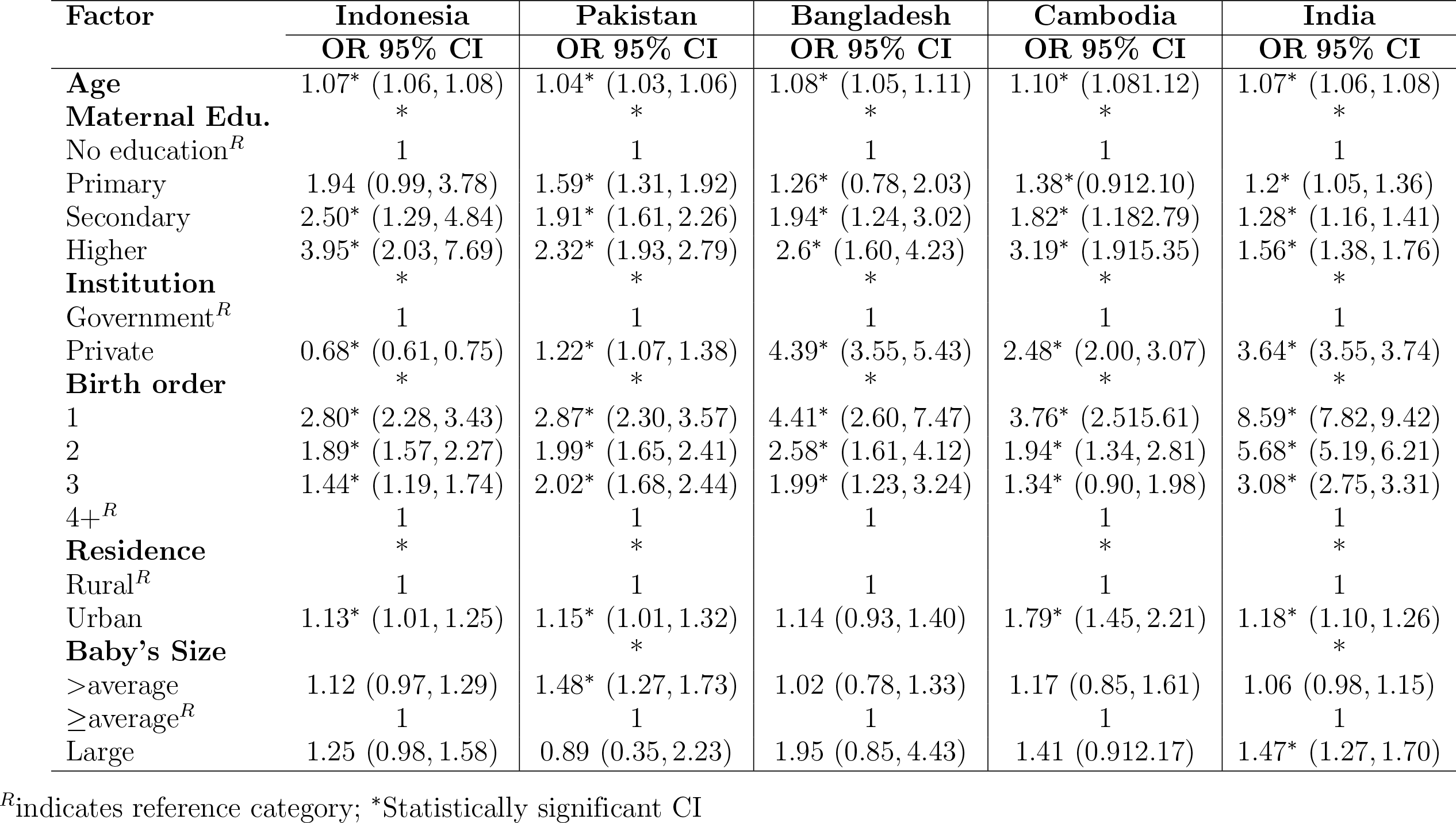
Adjusted odds ratio (OR) and 95% confidence interval (CI) for the risk of caesarean section corresponding to the associated factors in **South and South-east Asian Countries**

**Table 5.**
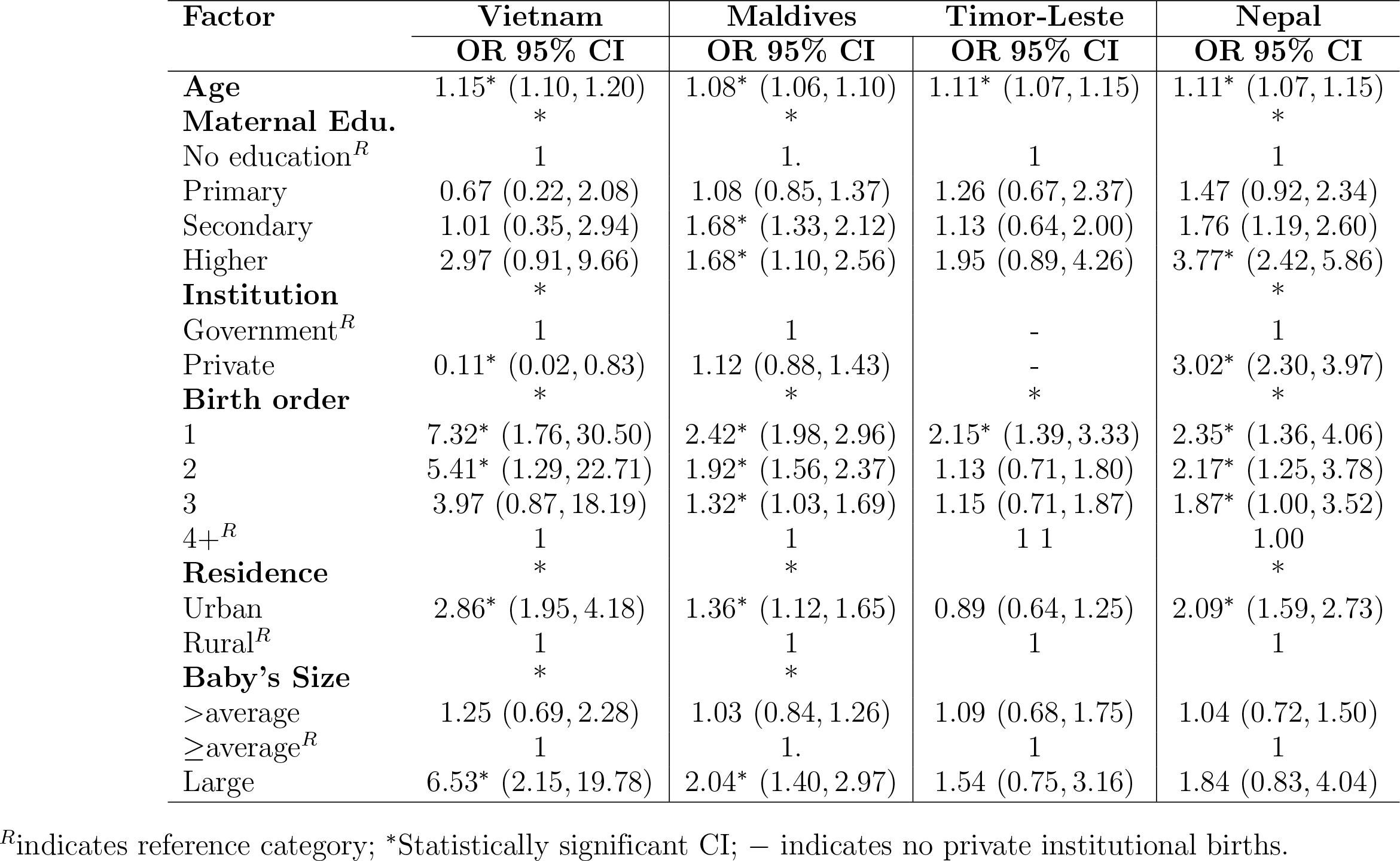
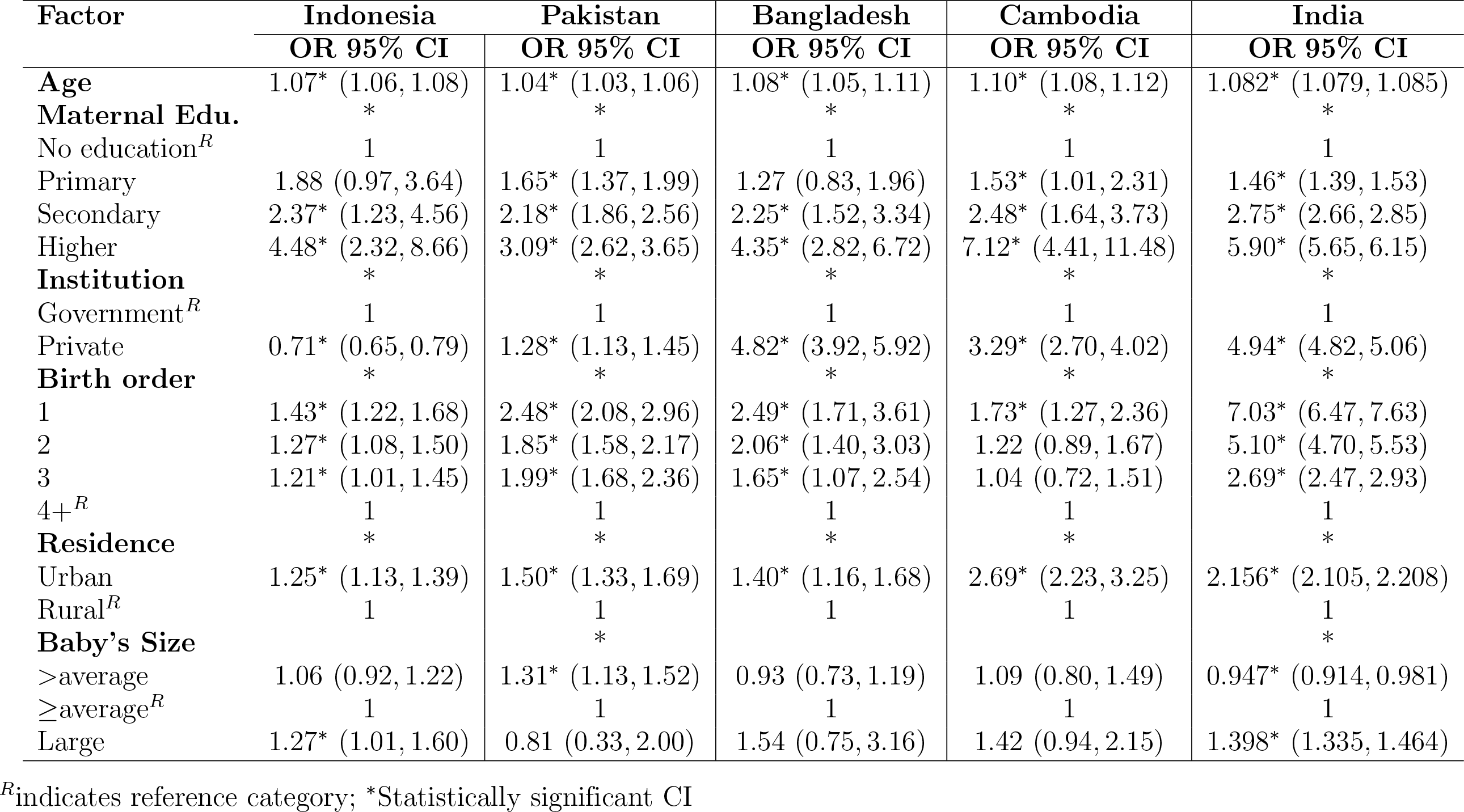
Unadjusted odds ratio (OR) and 95% confidence interval (CI) for the risk of caesarean section corresponding to the associated factors in **South and South-east Asian Countries**

| Factor | Vietnam | Maldives | Timor-Leste | Nepal |
| --- | --- | --- | --- | --- |
|  | OR 95% CI | OR 95% CI | OR 95% CI | OR 95% CI |
| **Age** | 1.15* (1.10, 1.20) | 1.08* (1.06, 1.10) | 1.11* (1.07, 1.15) | 1.11* (1.07, 1.15) |
| **Maternal Edu.** | * | * |  | * |
| No education<sup>R</sup> | 1 | 1. | 1 | 1 |
| Primary | 0.67 (0.22, 2.08) | 1.08 (0.85, 1.37) | 1.26 (0.67, 2.37) | 1.47 (0.92, 2.34) |
| Secondary | 1.01 (0.35, 2.94) | 1.68* (1.33, 2.12) | 1.13 (0.64, 2.00) | 1.76 (1.19, 2.60) |
| Higher | 2.97 (0.91, 9.66) | 1.68* (1.10, 2.56) | 1.95 (0.89, 4.26) | 3.77* (2.42, 5.86) |
| **Institution** | * |  |  | * |
| Government<sup>R</sup> | 1 | 1 | - | 1 |
| Private | 0.11* (0.02, 0.83) | 1.12 (0.88, 1.43) | - | 3.02* (2.30, 3.97) |
| **Birth order** | * | * | * | * |
| 1 | 7.32* (1.76, 30.50) | 2.42* (1.98, 2.96) | 2.15* (1.39, 3.33) | 2.35* (1.36, 4.06) |
| 2 | 5.41* (1.29, 22.71) | 1.92* (1.56, 2.37) | 1.13 (0.71, 1.80) | 2.17* (1.25, 3.78) |
| 3 | 3.97 (0.87, 18.19) | 1.32* (1.03, 1.69) | 1.15 (0.71, 1.87) | 1.87* (1.00, 3.52) |
| 4+<sup>R</sup> | 1 | 1 | 1 1 | 1.00 |
| **Residence** | * | * |  | * |
| Urban | 2.86* (1.95, 4.18) | 1.36* (1.12, 1.65) | 0.89 (0.64, 1.25) | 2.09* (1.59, 2.73) |
| Rural<sup>R</sup> | 1 | 1 | 1 | 1 |
| **Baby's Size** | * | * |  |  |
| >average | 1.25 (0.69, 2.28) | 1.03 (0.84, 1.26) | 1.09 (0.68, 1.75) | 1.04 (0.72, 1.50) |
| ≥average<sup>R</sup> | 1 | 1. | 1 | 1 |
| Large | 6.53* (2.15, 19.78) | 2.04* (1.40, 2.97) | 1.54 (0.75, 3.16) | 1.84 (0.83, 4.04) |

<sup>R</sup>indicates reference category; \*Statistically significant CI; – indicates no private institutional births.

**Table 5 (cont.).** Unadjusted odds ratio (OR) and 95% confidence interval (CI) for the risk of caesarean section corresponding to the associated factors in South and South-east Asian Countries
| Factor | Indonesia<br>OR 95% CI | Pakistan<br>OR 95% CI | Bangladesh<br>OR 95% CI | Cambodia<br>OR 95% CI | India<br>OR 95% CI |
| --- | --- | --- | --- | --- | --- |
| **Age** | 1.07* (1.06, 1.08) | 1.04* (1.03, 1.06) | 1.08* (1.05, 1.11) | 1.10* (1.08, 1.12) | 1.082* (1.079, 1.085) |
| **Maternal Edu.** | * | * | * | * | * |
| No education<sup>R</sup> | 1 | 1 | 1 | 1 | 1 |
| Primary | 1.88 (0.97, 3.64) | 1.65* (1.37, 1.99) | 1.27 (0.83, 1.96) | 1.53* (1.01, 2.31) | 1.46* (1.39, 1.53) |
| Secondary | 2.37* (1.23, 4.56) | 2.18* (1.86, 2.56) | 2.25* (1.52, 3.34) | 2.48* (1.64, 3.73) | 2.75* (2.66, 2.85) |
| Higher | 4.48* (2.32, 8.66) | 3.09* (2.62, 3.65) | 4.35* (2.82, 6.72) | 7.12* (4.41, 11.48) | 5.90* (5.65, 6.15) |
| **Institution** | * | * | * | * | * |
| Government<sup>R</sup> | 1 | 1 | 1 | 1 | 1 |
| Private | 0.71* (0.65, 0.79) | 1.28* (1.13, 1.45) | 4.82* (3.92, 5.92) | 3.29* (2.70, 4.02) | 4.94* (4.82, 5.06) |
| **Birth order** | * | * | * | * | * |
| 1 | 1.43* (1.22, 1.68) | 2.48* (2.08, 2.96) | 2.49* (1.71, 3.61) | 1.73* (1.27, 2.36) | 7.03* (6.47, 7.63) |
| 2 | 1.27* (1.08, 1.50) | 1.85* (1.58, 2.17) | 2.06* (1.40, 3.03) | 1.22 (0.89, 1.67) | 5.10* (4.70, 5.53) |
| 3 | 1.21* (1.01, 1.45) | 1.99* (1.68, 2.36) | 1.65* (1.07, 2.54) | 1.04 (0.72, 1.51) | 2.69* (2.47, 2.93) |
| 4+<sup>R</sup> | 1 | 1 | 1 | 1 | 1 |
| **Residence** | * | * | * | * | * |
| Urban | 1.25* (1.13, 1.39) | 1.50* (1.33, 1.69) | 1.40* (1.16, 1.68) | 2.69* (2.23, 3.25) | 2.156* (2.105, 2.208) |
| Rural<sup>R</sup> | 1 | 1 | 1 | 1 | 1 |
| **Baby's Size** |  | * |  |  | * |
| >average | 1.06 (0.92, 1.22) | 1.31* (1.13, 1.52) | 0.93 (0.73, 1.19) | 1.09 (0.80, 1.49) | 0.947* (0.914, 0.981) |
| ≥average<sup>R</sup> | 1 | 1 | 1 | 1 | 1 |
| Large | 1.27* (1.01, 1.60) | 0.81 (0.33, 2.00) | 1.54 (0.75, 3.16) | 1.42 (0.94, 2.15) | 1.398* (1.335, 1.464) |

<sup>R</sup>indicates reference category; \*Statistically significant CI

Women with higher education are more likely to have a caesarean section compared to those who have less education. Women who have opted for private institutions for delivery, compared to governmental medical institutions, are more likely to undergo a caesarean section. The order of birth showed a constant decrease in caesarean section. The place of residence showed that urban women, compared to their rural counterparts, have seen to be more likely to experience caesarean sections. The size of the baby has found significant in some countries except Timor-Leste, Nepal, Indonesia, Bangladesh and Cambodia, which shows that women in these countries whose baby size is above average or below average with reference to average size are more likely to give birth by caesarean section.

As caesarean section is a surgical procedure and is possible only if the location is equipped with medical facilities; therefore, women whose delivery is not institutional cannot be considered to be exposed to a caesarean delivery and are not part of the population of interest. Only women who were married and whose deliveries were institutional are considered to be within the population of interest. Corresponding to each selected country, the results revealed a shift towards institutional delivery over those of non-institutional deliveries, which indicates the effectiveness of health programs and mothers’ increasing awareness of the importance of an institutional delivery. It should also be noted regarding the Maldives that the prevalence of caesarean sections based on the total number of deliveries was 31.78%, which is quite close to the prevalence based on births in institutions, 33.14%. The reason for this closeness of estimates is that in the Maldives, 95.94% of births are institutional. In the case of Timor-Leste and Bangladesh, respectively, where 19.21% and 23.52% of births are in institutions. The prevalence based on the institutional births in Timor-Leste and Bangladesh has found to 4.45 and 5.2 times higher than those rates based on the total number of births, respectively. Our findings suggest that women with higher education are more likely to undergo caesarean as compared to uneducated women. The age of women was found to have a weak positive impact (the odds are slightly higher than one) on the risk of caesarean (i.e., every one-year increment on women’s age, the risk of caesarean is approximately 1.1 times in women as compared to women with normal delivery). There are positive trends found for caesarean delivery in private hospitals. Our results indicate that odds for caesarean delivery in private hospitals as compared to government hospitals are very high. Results suggest that women with higher education are more likely to have caesarean sections than women who are uneducated. The age of women with low impact (odds are slightly higher than 1) on the risk of caesarean section. Results have indicated that odds for caesarean delivery in private hospitals as compared to government hospitals are very high.

## Conclusions

There is a significant inclination in institutional deliveries in all South and Southeast Asian countries, which indicates the effectiveness of women’s awareness programs and programs. The main reason for this transition is that it reduces the risks and complications that occur during deliveries. This increase in the number of deliveries may be an important reason for caesarean delivery bias in all South and Southeast Asian countries (except for Vietnam, Timor-Leste, Nepal, and Cambodia) above the previous WHO recommended optimal range of 10-15% and the more recent recommendation of top researchers of 19%. Having said this, the recommended rate might be higher if preventing serious morbidity is also taken into account. The prevalence of caesarean section is also examined for different socioeconomic covariate markers. The analysis shows that maternal age, maternal education, and birth order are significantly associated with caesarean delivery.

Of all the other determinants of the prevalence of caesarean delivery in any medical facility, the choice of place of birth *viz.*, Government and private facilities may be a strong influence on the choice to undergo a caesarean section. Increases in the caesarean rates create a heavy burden on the health system [15] and also increases the risk of other health problems to both mother and baby, and unwanted caesarean delivery also puts a huge financial burden on the family economic status.

This study examines the prevalence of caesarean section in selected countries of South and Southeast Asia, which is the top and third largest populous region in the world. To represent the difference between rural and urban areas, the DHS datasets are classified so that the variation between the various facilities and the health and demographic indicators can be examined according to their place of residence. The study has also shown a behaviour change in rural and urban areas towards the adoption of a caesarean procedure. But in urban areas, rates are comparatively high in rural areas. The results showed that despite the disparity in the prevalence of caesarean section among rural and urban women, the percentages based on institutional births are completely different from those obtained using the information on total births. As a result, the government needs to develop better healthcare infrastructure and along with more antenatal care related schemes to reduce the risks associated with increased caesarean delivery.

## Acknowledgements

The authors would like to express their deepest gratitude and sincerely thank to the Demographic and Health Surveys (DHS) programme, to provide access to data for this study.

## Disclosure statement

No potential conflict of interest was reported by the authors.

